# Long-term, functional culture and *in vitro* manipulation of adult mouse cardiomyocytes

**DOI:** 10.1101/721514

**Authors:** Neal I. Callaghan, Shin-Haw Lee, Sina Hadipour-Lakmehsari, Xavier A. Lee, M. Ahsan Siraj, Amine Driouchi, Christopher M. Yip, Mansoor Husain, Craig A. Simmons, Anthony O. Gramolini

## Abstract

Primary adult cardiomyocyte (aCM) culture is challenged by poor survival and loss of phenotype, rendering extended *in vitro* experiments unfeasible. Here, we establish murine aCM culture methods that enhance survival and maintain sarcomeric structure and Ca^2+^ cycling to enable physiologically-relevant contractile force measurements. We also demonstrate genetic and small-molecule manipulations that probe mechanisms underlying myocyte functional performance.

---

*In vitro* primary cell culture is a valuable tool to complement *in vivo* physiological investigation of many tissues. *In vitro* studies of myocardium are limited by the challenges of adult cardiomyocyte (aCM) culture. Although neonatal and pluripotent stem cell-derived cardiomyocytes have high survival rates in culture, they do not fully replicate the adult phenotype in terms of morphology, mature protein isoform expression, action potential component currents, Ca^2+^ transient dynamics, contractile force production, or metabolic substrate preference^1,2^. Therefore, these immature cells fall short of replicating the physiology of mature CMs required for detailed mechanistic study^1^. While primary aCM isolation remains a crucial tool for acute studies^2^, primary aCMs cultured in single-cell format detach and rapidly lose physiological function^2–4^, with surviving cells typically assuming a ‘fetal’ phenotype after 2-3 weeks in culture^2^. Finally, aCM contractility has been primarily measured using shortening velocity of cells in suspension or isometric force production of intact or permeabilized cells glued to a force transducer. Neither of these methods represent a physiological setting. aCMs respond in real time to their physical environment^5,6^, and commonly used contractile metrics of shortening velocity of aCMs in suspension are not ideal because they do not provide physiological resistance nor estimate contractile force directly. Here, we demonstrate a culture protocol that maintains primary murine aCM function, enables physiological contractile force measurement, and is amenable to genetic and small-molecule treatments with phenotype-altering effects.

Typically, primary CMs are often cultured on a purified matrix and supplemented with specific growth factors added to the media. Since aCM function depends on various integrin-ECM protein interactions to allow for precise regulation of downstream signaling pathways^7,8^, we first explored altering the matrix constituents to promote aCMs. Basement membrane preparations including Geltrex provide a complex ECM protein mixture including collagens, laminins, fibronectins, and entactins among others^9^, recapitulating the diversity of composition of intact myocardial ECM^10^. Secondly, the strength of aCM contractions result in substantial stresses on the membranes of these cells. In aCM culture, butanedione monoxime (BDM) was initially adopted as a myosin ATPase inhibitor, however BDM also impairs Ca^2+^ cycling through L-type channels^11,12^, modulates cardiac ryanodine receptor (RyR) flux in a Ca^2+^-dependent manner^13^, and inhibits the oxidative metabolism upon which aCMs are reliant^14^; all of which are deleterious to maintaining cellular myocyte homeostasis^15^. A non-BDM protocol to inhibit contractility, using myosin ATPase inhibitor blebbistatin, has shown enhanced aCM longevity and function in culture^3^.

Previous methods attempting prolonged aCM survival have shown complete loss of initial sarcomere structure by day 8 (d8)^2^, followed by regressionto a neonatal morphology. However, CMs isolated from adult mice using our modified protocol combining Geltrex and blebbistatin demonstrated high adhesion, survivability and continued functionality in sustained culture (Figure 1). CMs retained α-actinin-positive sarcomeres and an ordered sarcoplasmic reticulum (SR) with both longitudinal and cisternal components (Fig. 1A). DHPR-positive t-tubules exhibited a mild but progressive degree of disorder starting even on d1 after isolation; overall sarcolemmal organization as evidenced by STIM1 imaging revealed a similar trend. Furthermore, in the presence of blebbistatin the use of Geltrex resulted in a 32 ± 11% decrease in Akt phosphorylation compared to a standard laminin coating (p = 0.02, Fig 1B), indicating potentially altered signaling in pathways associated with differentiation, proliferation, and growth. This was reflected in the survival and morphology of aCMs, which remained largely viable and only declined slowly from an initial yield of 90-95% rod-shaped cells (Fig. 1C); surviving aCMs exhibited morphology similar to their initial state immediately post-isolation for up to 1 week in culture and neither hypertrophied nor lost phenotype, as is common in standard aCM culture methods.

**Figure 1.**
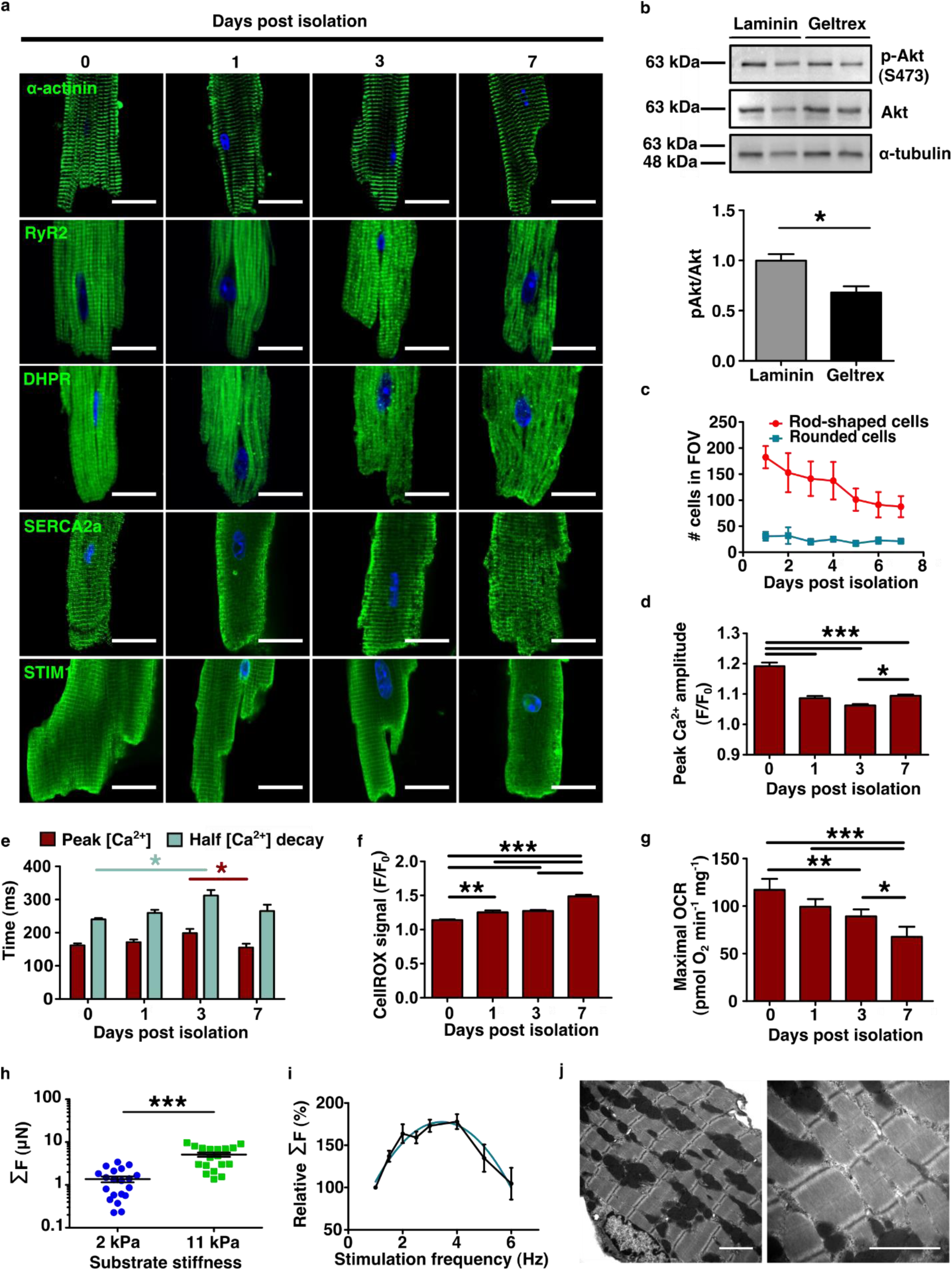
Isolated adult murine cardiomyocytes (aCMs) show sustained viability and retention of functional protein expression patterns up to 7 days post isolation (DPI). A) Retention of actinin-positive sarcomeres and calcium handling protein expression patterns (green) up to one week in culture with no evidence of hypertrophic remodeling or dedifferentiation. Scale bars equal 20 µm. Nuclei represented in blue with Hoechst 33342. B) Culture of aCMs for 24 h on Geltrex-coated surfaces induces lower Akt activation (S473-Pi) than parallel cultures plated on laminin-111 (N=3), p=0.023. C) Survival of viable rod-shaped (red) relative to rounded CMs (blue) (N=3). Evidence of calcium handling adaptation *in vitro* as peak calcium transient signal amplitude (D) normalized to baseline (F/F_0_) partially recovers by 7 days from a minimum at c.a. 3 DPI, as do E) rising time to peak amplitude and falling time from peak to half-peak amplitude. F) Normalized intracellular CellROX signal increases over time in culture (N=3). G) Maximal FCCP-uncoupled oxygen consumption rate (OCR) decreases with culture period while glycolytic extracellular acidification rate (ECAR) increases (N=3). H) Spontaneously-contracting CMs generate higher peak contractile force on a TFM substrate of 11 kPa stiffness than one of 2 kPa (N=20 for each treatment). I) Electrically-stimulated CMs show a physiological (Bowditch)-resembling force-frequency relationship that scales relative to the absolute force of the cell at *f*=1 Hz (N=5), quadratic fit R^2^ = 0.61. J) TEM reveals detailed sarcomeric substructure in isolated aCMs. All data expressed as mean ± SEM; N denotes biological replicates; significance indicated by * (p<0.05), ** (p<0.01), and *** (p<0.001).

Spontaneous Ca^2+^ transient properties were analyzed using Fluo-4 AM fluorescent microscopy (Figure 1D-E) after 1 week in culture. The magnitude of Ca^2+^ transients sharply decreased at d1 but remained relatively stable thereafter. Kinetics of the transients tended to slow after d1, becoming significantly slower by d3, and recovering to near-initial levels by d7. A decrease in Ca^2+^ transient magnitude, peaking at d3, coincided with the most prolonged transient rise and decay times. Cells at d7 showed a recovery in transient magnitude and decay time towards the initial measurements, suggesting acclimatization to culture. The decreased regularity in DHPR expression throughout the cell suggested that t-tubule organization was somewhat disrupted after extended culture. There was little evidence of SR disruption as a function of extended culture time, as expression patterns and levels of RyR2 and SERCA2a remained relatively constant. Since the L-type Ca^2+^ current is necessary to induce SR Ca^2+^ release, aberrant DHPR expression corresponding to disrupted t-tubules would likely result in lower velocity of both sarcolemmal and sarcoplasmic contributions to Ca^2+^ flux. Similar changes to STIM1 expression patterns over time suggest that membrane and t-tubule physiology continues to adapt *in vitro*. The rate of CellROX-active ROS production through live cell imaging increased steadily in culture to 149 ± 10% of baseline at d7 (p < 0.0001; Fig. 1F). This increase coincided with a significant decrease in FCCP-uncoupled maximal oxygen consumption rate (p < 0.0001, Fig. 1G), suggesting growing metabolic inefficiency over increased culture time. However, increased maximal to basal OCR ratio (4.3 vs. 3.0) was noted at d1 compared to standard Langendorff isolation and culture methods^16^, suggesting significant spare respirometric capacity early in culture with this protocol.

Traction force microscopy (TFM) has been carried out in many cell types, including neonatal and iPSC-derived cardiomyocytes ^17^. However, its use in primary isolated aCMs has been hampered by poor cell attachment and functionality post-isolation. In this study, due to the significant improvement of cell isolation, we applied widefield TFM techniques to aCMs to measure auxotonic contractions in a mechanically-relevant environment. Spontaneous cell-associated total stresses ranging from 1.1-6.1 kPa were used to calculate single-cell contractile forces of 1.4-9.6 µN cell^-1^ or estimated cross-sectional forces of 16.5 ± 2.5 mN mm^−2^ (mean ± SEM). These cross-sectional force values are in agreement with other auxotonic measurements of individual cells obtained with considerably higher experimental difficulty^18^. Cells plated on 2 kPa gels produced significantly lower peak forces than cells on gels of 11 kPa (Fig. 1H). When electrically paced, cells showed a positive force-frequency response, peaking between 2.5-4 Hz and 180% of baseline 1 Hz force (Fig. 1I) and approximating a quadratic curve. The resulting Bowditch curve was significantly steeper and left-shifted compared to *in vivo* measurements of mice^19^, ostensibly due to the effects of isolation and culture. Similarly, cells failed to reliably pace at field stimulation rates above 7 Hz while murine myocardium can pace *in vivo* to *c.a.* 14 Hz. Transmission electron microscopy (Fig. 1J) revealed preserved sarcomeric structures and mitochondrial networks.

Finally, we demonstrated functional manipulation of aCMs entirely *in vitro* (Figure 2). In phospholamban (PLN)-knockout CD1 mice we showed lentiviral transfection of WT-PLN and the human pathogenic variant R9C-PLN was successful (Fig. 2A), and clearly visualized FLAG-PLN expression (Fig. 2B). Furthermore, TFM analysis revealed differences between peak contractile forces, likely due to different levels of SERCA2a regulation between treatments (Fig. 2C). These experiments clearly highlight the feasibility of utilizing these aCMS for biochemical and functional assays in genetic studies. In parallel experiments, we showed that we can knockdown gene expression in these cells, further emphasising the utility of the cellular isolation methodology. Specifically, as the SR adaptor protein Reep5 is known to be essential in maintaining cardiac function^20^, AAV9-mediated *Reep5* shRNA knockdown (Fig. 2D) induced disorganized SR/ER morphology (Fig. 2E) and increased spontaneous Ca^2+^ efflux time relative to treatment with scrambled control shRNA (Fig. 2F). Similarly, ER stress induced by 24 h tunicamycin treatment (5 µmol L^-1^) resulted in extensive t-tubular disorganization and vacuolization as assessed by dSTORM super-resolution imaging (Fig. 2G; analysis in Fig. 3). These sample workflows can be adapted for *in vitro* characterization of pathways and mechanisms of interest in early disease progression, similar to those already performed in primary cultures of other tissues.

**Figure 2.**
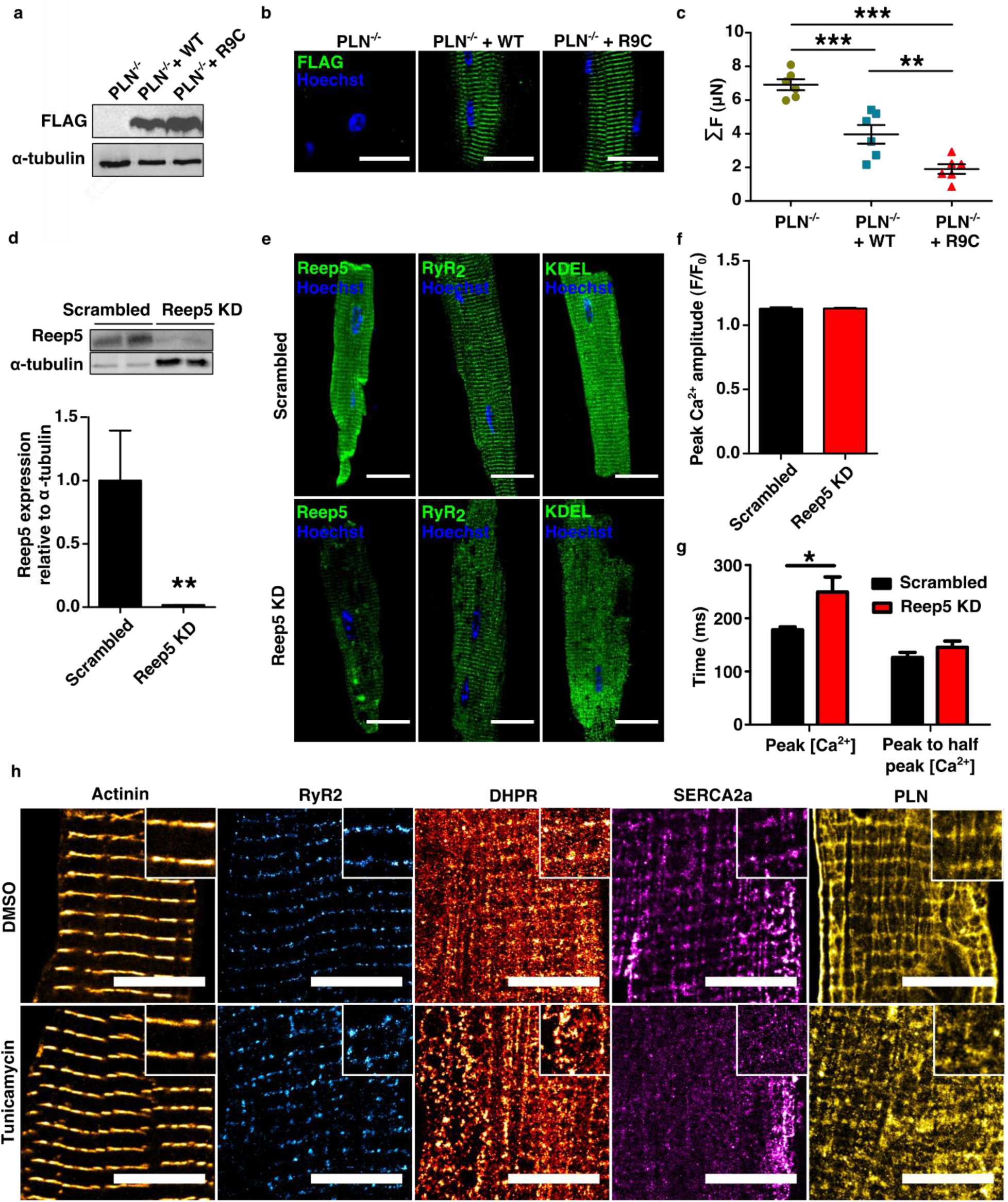
Isolated murine aCMs can be functionally modulated by *in vitro* treatments. A) WT and human pathogenic R9C phospholamban (PLN) cDNA can be expressed in sarcomeres using a lentiviral vector *in vitro* from aCMs isolated from a PLN-null mouse; scale bars represent 20 µm. B) Confirmation of PLN-FLAG expression post lentiviral transfection. C) Traction force microscopy reveals contractile differences corresponding to the different levels of activity of phospholamban variants. D) Reep5 shRNA-mediated knockdown by AAV9 transduction is confirmed by immunoblot (Reep5 KD treatment loaded at higher volume to produce quantifiable bands). E) Confocal images confirm knockdown of *Reep5* and resulting disorganization of SR (RyR2) and ER (KDEL motif) compared to scrambled control; scale bars represent 20 µm. F) Peak calcium transient amplitude is not significantly affected by *Reep5* KD (N=6). G) Time to peak calcium transient amplitude is significantly higher after *Reep5* KD, but transient decay time to 50% is not affected. H) Super-resolution (dSTORM) microscopy reveals that aCMs are responsive to tunicamycin-induced protein folding stress (5 µmol L^-1^ for 24 h) as seen by the disruption of calcium-handling protein expression patterns. Scale bars represent 10 µm, magnified inset images are 4×4 µm. All data expressed as mean ± SEM; N denotes biological replicates; significance indicated by * (p<0.05), ** (p<0.01), and *** (p<0.001).

**Figure 3.**
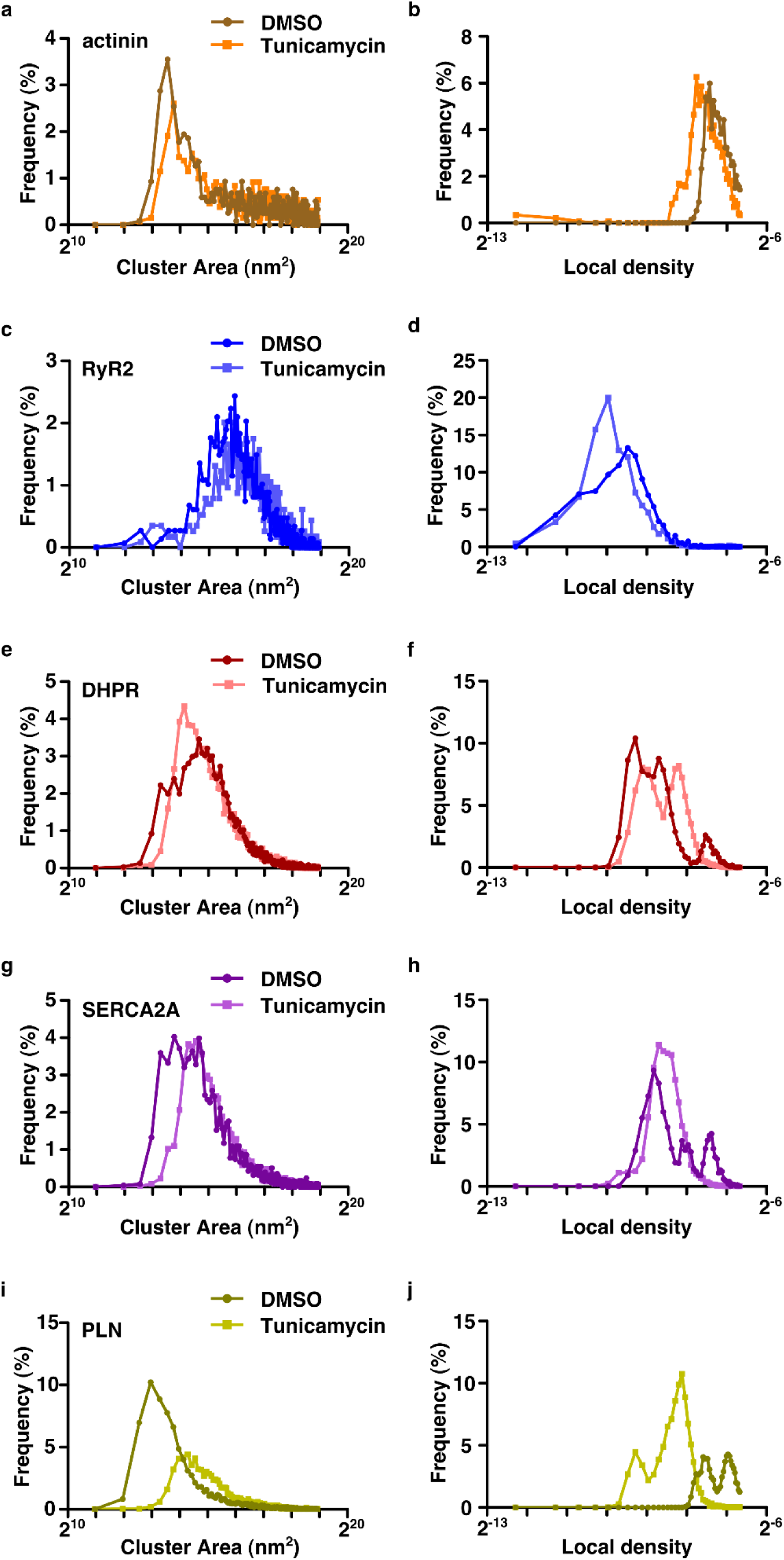
Voronoi tessellation cluster analysis of dSTORM-imaged aCMs in Figure 2. Staining for actinin (A-B), RyR2 (C-D), DHPR (E-F), SERCA2A (G-H), and PLN (I-J). Cluster area (A, C, E, G, I) and cluster density (B, D, F, H, J) was analyzed after 24 h treatments with either a DMSO sham or tunicamycin (5 µmol L^-1^).

In conclusion, this study demonstrates a significant innovation to primary murine aCM cultures with increased cell longevity and durable physiological function *in vitro*, notably allowing non-acute experimental manipulations in aCM culture for the first time. By combining the physiological benefits of blebbistatin and Geltrex, the loss of aCM phenotypes and cell death is reduced, with aCM hallmarks retained over at least 7 days of culture. Importantly, we have demonstrated the ability to model pathology through both genetic modulation and small-molecule administration due to the enhanced timescales and experimental flexibility afforded by these methods. Finally, to our knowledge, we also describe the first use of live-cell TFM for physiological, treatment-sensitive force measurement of primary aCM auxotonic contraction. Together, these tools will allow for the application of experimental treatments and measurements previously restricted to *in vivo* settings to an *in vitro* model, with the mechanistic control afforded by the single-cell scale.

## Methods

### Ethical statement

All studies were approved by the University of Toronto Animal Care Committee and conducted under its Animal Care Guidelines.

### Reagents, media, and consumables

Perfusion and EDTA buffers were prepared as previously described^2^, except that they contained 15 µmol L^-1^ (S)-(-)-blebbistatin (Toronto Research Chemicals, Toronto, ON). EDTA, taurine, and HEPES were supplied by BioShop Inc. (Burlington, ON). M199 (Wisent Inc., Saint-Jean-Baptiste, QC) pH 7.6 was supplemented with added 100X CD lipid, 100X insulin-transferrin-selenium supplement (Life Technologies), and 100X penicillin/streptomycin to 1X each. All other reagents unless mentioned were supplied by Sigma-Aldrich (St. Louis, MI).

### Isolation of primary murine adult cardiomyocytes

The surgical and perfusion methods closely followed those of a recent optimization of a Langendorff-free preparation^2^; we encourage readers to refer to Ackers-Johnson *et al.* for detailed considerations in surgical methods and culture optimization. We have provided media recipes in Supplementary Table 1. Briefly, male CD1 mice of 8 weeks or older were euthanized by open drop exposure to isoflurane followed by cervical transection. The chest cavity was opened, the descending aorta severed, and 7 mL of EDTA buffer containing 15 µmol L^-1^ blebbistatin (Toronto Research Chemicals, Toronto ON) was injected into the right ventricle. The heart was hemostatically clamped at the ascending aorta, excised from the chest cavity, and placed into a fresh dish of EDTA buffer with blebbistatin while 9 mL of the same buffer was slowly injected into the apex of the left ventricle. After the heart was free of blood, it was moved to a dish of perfusion buffer with 15 µmol L^-1^ blebbistatin, and injected with 3 mL of fresh 15 µmol L^-1^ blebbistatin through the same hole previously used in the LV. Finally, the heart was moved to a dish containing 475 U mL^-1^ collagenase type II (Worthington Biochemical Corporation, Lakewood NJ) in perfusion buffer with 15 µmol L^-1^ blebbistatin, of which 20 mL more was injected through the existing LV hole. We found that the use of 27G, ½ length needles minimized mechanical damage to the heart, allowing for maintained pressure during perfusion and optimal coronary circulation of collagenase and thus digestion of the myocardium. Additionally, we observed batches of higher specific activity (IU/mg) collagenase type II to be more amenable to cell survival, even at the same final activity concentration.

**Table 1.** Composition of EDTA buffer, perfusion buffer, cell culture medium, and plating medium. All buffers are made in >18 M? deionized water, sterile-filtered, and replaced frequently.

| **Buffer** | **Molar mass (g mol<sup>-1</sup>) or stock concentration** | **Final concentration** | **Amount per 100 mL** |
| --- | --- | --- | --- |
| **EDTA buffer (pH 7.80)** |  |  |  |
| NaCl | 58.44 | 130 mmol L<sup>-1</sup> | 759.7 mg |
| KCl | 74.55 | 5 mmol L<sup>-1</sup> | 37.3 mg |
| NaH<sub>2</sub>PO<sub>4</sub> | 119.98 | 0.5 mmol L<sup>-1</sup> | 6.0 mg |
| HEPES | 238.3 | 10 mmol L<sup>-1</sup> | 238.3 mg |
| Glucose | 180.16 | 10 mmol L<sup>-1</sup> | 180.2 mg |
| Taurine | 125.15 | 10 mmol L<sup>-1</sup> | 125.2 mg |
| EDTA | 292.24 | 5 mmol L<sup>-1</sup> | 146.1 mg |
| Blebbistatin* | 292.33 | 0.015 mmol L<sup>-1</sup> | 438 µg |
| **Perfusion Buffer (pH 7.80)** |  |  |  |
| NaCl | 58.44 | 130 mmol L<sup>-1</sup> | 759.7 mg |
| KCl | 74.55 | 5 mmol L<sup>-1</sup> | 37.3 mg |
| NaH<sub>2</sub>PO<sub>4</sub> | 119.98 | 0.5 mmol L<sup>-1</sup> | 6.0 mg |
| HEPES | 238.3 | 10 mmol L<sup>-1</sup> | 238.3 mg |
| Glucose | 180.16 | 10 mmol L<sup>-1</sup> | 180.2 mg |
| Taurine | 125.15 | 10 mmol L<sup>-1</sup> | 125.2 mg |
| MgCl<sub>2</sub> | 95.21 | 1 mmol L<sup>-1</sup> | 9.5 mg |
| Blebbistatin* | 292.33 | 0.015 mmol L<sup>-1</sup> | 438 µg |
| Collagenase type II† | - | 475 U mL<sup>-1</sup> | 47500 U |
| FBS‡ | - | 10% | 10 mL |
| **Culture medium (pH 7.60)\*\*** |  |  |  |
| M199<br>(with NaHCO<sub>3</sub>, HEPES, L-glutamine, and 5 mM glucose) | - | - | 97 mL |
| Chemically defined lipid supplement | 100X | 1X | 1 mL |
| Insulin-transferrin-selenium supplement | 100X | 1X | 1 mL |
| Penicillin/Streptomycin | 100X | 1X | 1 mL |
| Blebbistatin* | 292.33 | 0.015 mmol L<sup>-1</sup> | 438 µg |
| FBS‡‡ | - | 5% | 5 mL |

\*Blebbistatin should be added fresh daily from a dry or DMSO frozen stock. Do not expose to light.
†Add collagenase type II only to create fresh collagenase buffer.
‡Add FBS without collagenase to create stop buffer. Make fresh daily.
\*\*Bicarbonate-free, blebbistatin-free cell culture medium can be used for contractile analysis. Cells should begin beating <5 min after medium change.
‡‡Add FBS to create plating medium. Adjust other volumes accordingly. Make fresh daily.

At this point, the tissue was minced in 3 mL of fresh collagenase buffer with two pairs of forceps, and gently triturated with a wide-bore 1 mL pipette. The collagenase activity was inhibited with addition of 3 mL of perfusion buffer with blebbistatin and 10% FBS (Wisent)). The isolate was then passed through a 70 µm strainer and rinsed with 3 mL additional stop buffer. The filtrate was divided between two 15 mL Falcon tubes which were left standing upright for 15 min. The rod-shaped viable cardiomyocytes gravity-settled to form a deep red pellet, while rounded, nonviable CMs and other cell types remained in suspension. The use of 2 tubes prevented oxygen or nutrient gradients forming in the cell pellets, while the use of steep-walled 15 mL Falcon tubes allowed for the best recovery of the pellet over successive washes. The supernatant was carefully removed, and the cells resuspended in a mixture of 75% perfusion buffer and 25% culture media, containing 15 µmol L^-1^ blebbistatin. Cells were allowed to settle 15 min, and the process repeated two more times with mixtures of 50%:50% and 25%:75% perfusion buffer:culture media respectively, all containing 15 µmol L^-1^ blebbistatin. The final cell pellet was resuspended in culture medium containing 5% FBS and 15 µmol L^-1^ blebbistatin, which was then plated for culture in the required format. After having been plated for 3 h, dishes were gently washed and replaced with culture medium containing 15 µmol L^-1^ blebbistatin (no FBS) to avoid serum toxicity. Cells were then cultured typically up to 7 DPI for functional analyses, although we noted substantial CM survival 3 weeks post-isolation without full phenotypic characterization.

### Cell culture and treatments

Cells were cultured on 14 mm glass coverslips in 35 mm dishes (P35GCOL-1.5-14-C, MatTek, Ashland, MA) coated with 250 µL Geltrex diluted 1:100 in M199 for immunofluorescence, ROS, and Ca^2+^ imaging at ∼100,000 cells/coverslip. For immunoblotting, cells were plated in 6-well tissue culture polystyrene (TCPS) plates at c.a. 1×10^6^ cells/well. For Agilent Seahorse respirometric characterization, cells were plated at ∼ 250,000 cells well^-1^ in 24-well Seahorse plates coated with 50 µL Geltrex diluted 1:100 in M199. 6-well surfaces were coated with 1.5 mL Geltrex diluted 1:100 in M199, except for laminin-111 (Trevigen Inc., Gaithersburg, MD, USA) at 5 µg mL^-1^ for direct Geltrex-laminin comparisons.

For viral transfection, aCMs 4 h post-plating, pretreated with 10 µg mL^-1^ polybrene (TR-1003, Millipore-Sigma), were treated for a further 21 h with a lentiviral vector containing PLN-WT or PLN-R9C, or an AAV9 vector containing Reep5 or scrambled shRNA, prepared as previously described^21^. For the PLN experiment, cells were fixed or subjected to TFM analysis 24 h after viral introduction. For the Reep5 experiment, cells were incubated a further 24 h after the vector was removed before live Ca^2+^ imaging or fixation. For tunicamycin treatments, cells 4 h post-plating were incubated 24 h in 5 µmol ^L−1^ tunicamycin from *Streptomyces* sp. (T7765, Millipore-Sigma) from a 1 mg mL^-1^ DMSO stock and compared to a vehicle sham.

### Antibodies

Rabbit polyclonal anti-Akt antibody (1:1000 for IB, 9272; Cell Signaling Technology), rabbit polyclonal anti-pAkt-Ser473 antibody (1:1000 for IB, 9271; Cell Signaling Technology), rabbit polyclonal anti-α-tubulin antibody (1:1000 for IB, 2144, Cell Signaling Technology), mouse monoclonal anti-α-actinin antibody (1:400 for IF, A7811; Sigma-Aldrich), mouse monoclonal anti-ryanodine receptor antibody (1:100 for IF, ab2827; Abcam), mouse monoclonal anti-dihydropyridine receptor (DHPR) antibody (1:800 for IF, ab2864; Abcam), mouse monoclonal anti-sarco(endo)plasmic reticulum Ca^2+^-ATPase (SERCA) 2a antibody (1:200 for IF, MA3-919; Thermo-Fisher), rabbit polyclonal anti-STIM1 antibody (1:200, PA1-46217; Thermo-Fisher), mouse monoclonal PLN [2D12] antibody (1:500 for IF, 1:1000 for WB; ab2865, Abcam), mouse monoclonal FLAG antibody (1:500 for IF, 1:1000 for WB, F1804; Millipore-Sigma), mouse monoclonal anti-Reep5 antibody (1:500 for IF, 1:1000 for WB, 14643-1-AP; Proteintech), and rabbit monoclonal anti-KDEL antibody (1:250 for IF, ab176333; Abcam) were used in this study. Goat anti-rabbit Alexa Fluor 488 secondary antibodies (nos. A-11034 and A-11011; Molecular Probes) were used at 1:800 dilution.

### Immunoblotting

For Akt signaling analysis and genetic experiments, protein lysates from CMs confluently plated in a single well of a 6-well TCPS plate (0.5-1 heart well^-1^) were harvested in radioimmunoprecipitation assay buffer (RIPA, 50 mM Tris-HCl; pH7.4, 1% NP-40, 0.5% sodium deoxycholate, 0.1% SDS, 150 mM NaCl, 2 mM EDTA, 1X cOmplete Mini protease inhibitor cocktail (4693159001, Roche). For PLN expression analysis, cells were lysed in lysis buffer (8 mol L^-1^ urea, 10% (v/v) glycerol, 20% (w/v) SDS, 1 mol L^-1^ dithiothreitol, 1.5 mol L^-1^ Tris-HCl, pH 6.8, 1X cOmplete™ Mini protease inhibitor cocktail (4693159001, Roche)) with an 18-gauge needle. Lysates were centrifuged at 15,000 g for 15 min at 4°C.

SDS-soluble supernatants were added to 2X loading buffer and subjected to SDS-PAGE in a 12% polyacrylamide gel with 6% stacking gel at 100 V for 20 min, then 120 V for 1 h. Semi-dry transfer to a PVDF membrane occurred at 70 V for 1 h. Membranes were blocked in 5% BSA in TBS + 0.05% Tween-20 for 1 h at room temperature, then incubated overnight at 4°C in primary anti-Akt, anti-pAkt (Ser473), anti-Reep5, anti-FLAG, or anti-α-tubulin antibodies (described above), then in secondary antibodies (1:2500 dilution) for 1 h at room temperature. ECL detection was performed with a ChemiDoc™ Touch (Bio-Rad Laboratories, Hercules CA). Uncropped images are provided in Figure S1

### Confocal microscopy

Cultured cells were fixed with 4% paraformaldehyde for 10 min on ice, followed by 90% ice-cold methanol for 10 min. Next, cells were incubated with permeabilization buffer (0.5% Triton X-100, 0.2% Tween-20 in PBS) for 30 minutes at 4 degrees. Blocking buffer (5% FBS in 0.1% Triton-X-100 in PBS) was then added and incubated for 30 minutes at room temperature. Cells were incubated with primary antibodies (listed above) in blocking buffer (SERCA2a – 1:500, PLN – 1:1000, RyR2 – 1:1000, DHPR – 1:700) overnight at 4°C, and fluorophore-conjugated secondary antibody staining (Alexa 488; Molecular Probes) was performed at room temperature for 1 h in the dark. Nuclear counterstaining was performed using 1 μg/ml Hoechst 33342 (no. 4082; Cell Signaling) at room temperature for 15 min in the dark. Cells were imaged using a Zeiss spinning-disk confocal microscope.

### Traction force microscopy

TFM analysis in many cell types is often conducted by confocal microscopy, where a detergent is used to solubilize a cell to relieve its traction stress on gel. There, confocal microscopy allows for the imaging of only a single layer of gel. However, to characterize physiological CM contractions, widefield fluorescent microscopy is needed for temporal resolution. Therefore, TFM beads must be limited only to the surface of the gel. To this end, 18 mm circular coverslips were coated in a suspension of 500 nm red carboxylated FluoSpheres (580 nm excitation and 605 nm emission maxima; F8812, Thermo-Fisher) diluted 1:300 (v:v) in 100% ethanol, and slowly dried in a closed 12-well polystyrene culture plate to prevent heterogenous deposition of fluospheres. Matching coverslips were washed in 1 mol L^-1^ NaOH, washed with deionized water, dried, coated with (3-aminopropyl) triethoxysilane (APTES) for 10 min, then rinsed in deionized water again. An 11 kPa polyacrylamide (PA) solution (715 µL 50 mM HEPES pH 7.4, 150 µL 2% bis-acrylamide (Bio-Rad), 125 µL 40% acrylamide, 5 µL 10% ammonium persulfate, and 1 µL TEMED) was prepared, and 80 µL immediately spread on the silanized coverslip. 2 kPa PA gels were prepared similarly, except the volumes of HEPES, 2% bis-acrylamide, and acrylamide were 815, 40, and 137.5 µL respectively. The coverslip prepared with FluoSpheres was then gently floated on the solution, and the solution left to polymerize 30 min. The top coverslip was gently removed, leaving behind its FluoSphere coating at the surface of the gel. The gel remained conjugated to the APTES-functionalized bottom coverslip, which was placed in one well of a 12-well plate. Gels were washed 3x with PBS. Protein conjugation to the surface of the gel was accomplished as previously described ^22^. Briefly, N-sulfosuccinimidyl-6-(4’-azido-2’-nitrophenylamino) hexanoate (sulfo-SANPAH, CovaChem, Loves Park, IL) was solubilized in DMSO (0.25% final concentration) before diluting to 500 mmol L^-1^ in 50 mmol L^-1^ HEPES pH 7.4. The solution was immediately added (2 mL) to the well containing the gel, and exposed to 365 nm UV light for 10 min. This process was repeated with a fresh aliquot of sulfo-SANPAH solution. Gels were rinsed three times with 50 mM HEPES. Gels were then incubated overnight at 4°C in 1 mL 1:50 Geltrex in PBS. Gels were rinsed 3x with PBS before being plated with cells as previously described.

CMs were plated on the gels as described above. For contractile analysis, wells were rinsed and then replaced with blebbistatin-free culture media and incubated 5 min at 37 °C. Spontaneous contractions as visualized by displacement of the FluoSpheres were then recorded on an IX71 inverted widefield fluorescent microscope (Olympus Corporation, Tokyo, Japan) with a Texas Red filter cube at timelapse series with exposures of 55 ms. Brightfield images of the contracted cell were also taken for integration of the strain vectors (described below). For force-frequency curves, cells on 11 kPa gels were field-stimulated using two carbon electrodes of c.a. 1.5 cm length and 1 cm distance, soldered to copper leads that were then insulated with silicone rubber. For force-frequency curves, an S48 physiological monophasic square wave stimulator (Grass Technologies, Warwick, RI) was used to pace contraction from 1-6 Hz at 50 V, 5 ms duration.

Frames of peak contraction and relaxation were analyzed using a particle image velocimetry (PIV) plugin for ImageJ (v1.51j8)^23^. Interrogation windows of 64 × 64 pixels in 128 × 128 pixel search windows with a 0.60 correlation threshold were used to generate a displacement field, which was then used as the basis for Fourier transform traction cytometry (FTTC) as calculated by a separate ImageJ plugin ^23^. A Poisson’s ratio of 0.48, Young’s modulus of 11.0 kPa or 2.0 kPa, and a unitless regularization parameter (λ) of 4.7 × 10^−10^). The resulting stress matrix was integrated within the borders of the cell as captured by brightfield microscopy using a custom Matlab 2018a script. This sum was multiplied by the area of the cell to produce a total force scalar, and divided by 2 assuming both uniaxial force production by the cell, and null net force production given a complete reversion to the pre-contraction state. Final force values were expressed in terms of whole-cell peak force. Representative cells across N = 5 animals were measured for the 2 kPa vs. 11 kPa comparison and Bowditch curve. Duplicate separately-treated wells (one cell per well) for N = 3 animals were assessed for each treatment of the PLN-knockout experiment. TFM workflow is visualized in Figure 4.

**Figure 4.**
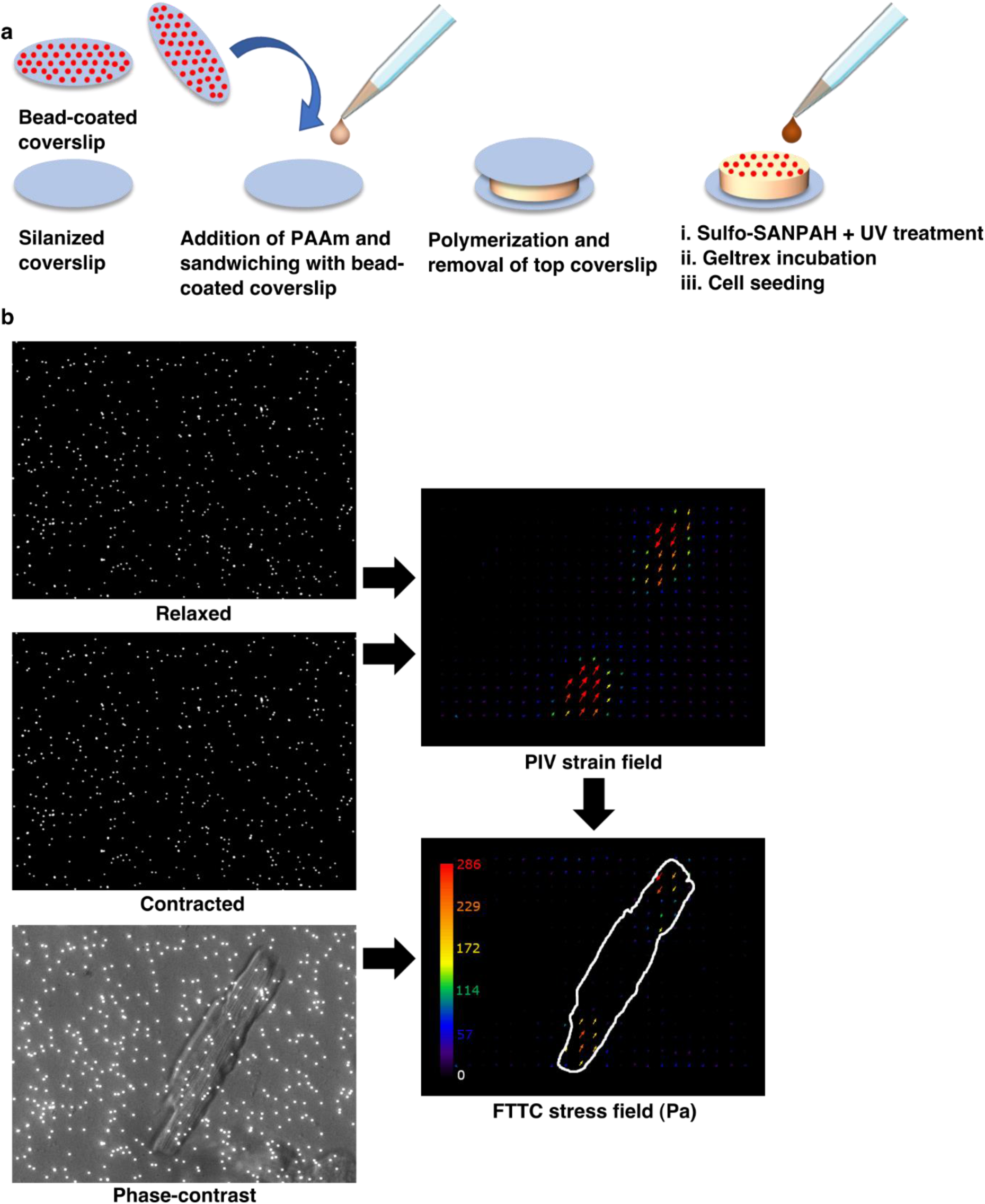
Sample TFM workflow demonstrating gel fabrication, stack acquisition, and data analysis. A) Preparation of a widefield-compatible TFM gel using 500 nm fluorescent microbeads. B) A strain field was constructed from frames of relaxation and maximal contraction before translation to a stress field using a Young’s modulus of 11 kPa for the polyacrylamide gel. The resulting uniaxial stress vectors were integrated within the projected area of the cell and divided by 2 to obtain a whole-cell net traction force.

### Reactive oxygen species (ROS) staining

CMs were incubated 30 min in culture medium with blebbistatin, containing 5 µmol L^-1^ CellROX Green (Thermo-Fisher) and counterstained 5 min with 1:1000 Hoechst 33342, both according to manufacturer directions. Cells were washed in fresh media and immediately imaged with a 40X objective on an inverted IX71 widefield microscope and using MicroManager acquisition software with exposures of 200 ms with a FITC filter and 50 ms with a DAPI filter. Fluorescent intensity per unit area was normalized to an equivalent area of adjacent background using ImageJ.

### Respirometry

Freshly isolated CMs were plated in 24-well Seahorse XFe™ (Agilent Technologies Inc, Santa Clara, CA, USA) plates and cultured as described previously for timepoint analysis. Mitochondrial respiration was assessed 0, 1, 3, and 7 DPI in a Seahorse XFe24 bioanalyzer. Cells were incubated in DMEM XF assay media (#102353-100, Agilent) supplemented with 5 mmol L^-1^ glucose, 1 mmol L^-1^ pyruvate, and 2 mmol L^-1^ glutamine at 37°C in a CO_2_-free incubator for 1 h prior to assay. Injector ports were loaded to provide final concentrations of 1 µmol L^-1^ oligomycin, 0.5 µmol L^-1^ FCCP, and 1 µmol L^-1^ rotenone and 2 µmol L^-1^ antimycin-A together, respectively according to the mitochondrial stress test protocol provided by Agilent. Maximal respiration was calculated and normalized to total protein concentration as measured by a Bradford assay. Optimal oligomycin and FCCP concentrations were previously determined by titration as shown in Supplemental Figure 3.

### Ca^2+^ imaging

Fluo-4 AM (Thermo-Fisher Scientific), reconstituted in DMSO and frozen at −20 °C, was incubated in the dark with cells at 37 °C for 30 min at a final concentration of 5 µmol L^-1^. Spontaneous Ca^2+^ waves were recorded using an IX71 widefield microscope and MicroManager acquisition software at 55 ms exposures with a FITC filter and expressed normalized to baseline fluorescence (F/F_0_) using ImageJ processing.

### Transmission electron microscopy

Isolated adult mouse cardiomyocytes were fixed in 2.5% glutaraldehyde in 0.1 mol L^-1^ phosphate buffer at 4°C overnight. Samples were post-fixed in 1% osmium tetroxide buffer and processed through graded alcohols and embedded in Quetol-Spurr resin. Sections of 90-100 nm were cut and stained with uranyl acetate and lead citrate and imaged at 20 000X magnification using a Hitachi TE microscope at the Department of Pathology, St. Michael’s Hospital (Toronto, Canada).

### dSTORM super-resolution imaging

Direct stochastic optical reconstruction microscopy was carried out as previously described ^24^. Briefly, stochastic photoswitching of immunostained samples was initiated with a buffer containing 50 mmol L^-1^ 2-mercaptoethylamine (M9768, Sigma-Aldrich), 40 µg mL^-1^ catalase (C3155, Sigma-Aldrich), 500 µg mL^-1^ glucose oxidase (G7141, Sigma-Aldrich), 50% (w/v) D-glucose (Sigma-Aldrich), in PBS pH 7.4. A 643 nm laser set at 20 mW was used to inactivate the AlexaFluor 647 fluorophore into an off-state prior to stochastic reactivation over the acquisition period. A super-resolved composite of 10 000 images acquired over a period of 300 s at 30 ms exposures was then reconstructed using the ThunderSTORM v1.3 ImageJ plugin, using a linear least square localization method. Coordinates of single emitters were filtered based on localization precision and photon count to discard electronic noise (0 nm < localization precision < 7 nm) and sample noise (localization precision >60 nm).

### Statistical analysis

All experiments were replicated at least 3 times. Statistical analysis was conducted with Prism 5 (GraphPad Software Inc.), except for respirometric analyses by JMP 11 (SAS Institute, Caly, NC, USA). All treatments were tested using the D’Agostino and Pearson omnibus normality test before comparison with an unpaired t-test. Respirometric timepoint data was assessed by 1-way split-plot ANOVA, followed by Tukey-Kramer HSD. Differences between timepoints and PLN cDNA knock-ins were assessed by 1-way ANOVA followed by Tukey’s LSD, except for Seahorse experiments which were assessed by 1-way repeated measure ANOVA followed by Tukey’s LSD. The contractile Bowditch curve was fitted with a quadratic regression (a = −11.91, b = 81.80, c = 37.08, R^2^ = 0.605). Differences were considered significant at p < 0.05.

## Acknowledgements

The authors thank Dr. Eric Strohm and Dorrin Zarrin-Khat for experimental assistance. The custom Matlab script was written by Richard Tam. PLN^−/−^ mice were a kind gift from Dr Evangelia Kranias (University of Cincinnati).

## References

1. Bedada, F. B., Wheelwright, M. & Metzger, J. M. Maturation status of sarcomere structure and function in human iPSC-derived cardiac myocytes. Biochim. Biophys. Acta - Mol. Cell Res. 1863, 1829–1838 (2016).

2. Ackers-Johnson, M. et al. A Simplified, Langendorff-Free Method for Concomitant Isolation of Viable Cardiac Myocytes and Nonmyocytes From the Adult Mouse Heart. Circ. Res. 119, 909–920 (2016).

3. Kabaeva, Z., Zhao, M. & Michele, D. E. Blebbistatin extends culture life of adult mouse cardiac myocytes and allows efficient and stable transgene expression. AJP Hear. Circ. Physiol. 294, H1667–H1674 (2008).

4. Banyasz, T. et al. Transformation of adult rat cardiac myocytes in primary culture. Exp. Physiol. 93, 370–382 (2008).

5. Helmes, M. et al. Mimicking the cardiac cycle in intact cardiomyocytes using diastolic and systolic force clamps; measuring power output. Cardiovasc. Res. 111, 66–73 (2016).

6. Pandey, P. et al. Cardiomyocytes Sense Matrix Rigidity through a Combination of Muscle and Non-muscle Myosin Contractions. Dev. Cell 44, 326-336.e3 (2018).

7. Schwartz, M. A. & Assoian, R. K. Integrins and cell proliferation: regulation of cyclin-dependent kinases via cytoplasmic signaling pathways. J. Cell Sci. 114, 2553–2560 (2001).

8. Harston, R. K. & Kuppuswamy, D. Integrins Are the Necessary Links to Hypertrophic Growth in Cardiomyocytes. J. Signal Transduct. 2011, 1–8 (2011).

9. Hughes, C. S., Postovit, L. M. & Lajoie, G. A. Matrigel: a complex protein mixture required for optimal growth of cell culture. Proteomics 10, 1886–1890 (2010).

10. Guyette, J. P. et al. Bioengineering Human Myocardium on Native Extracellular Matrix. Circ. Res. 118, 56–72 (2016).

11. Borlak, J. & Zwadlo, C. The myosin ATPase inhibitor 2,3-butanedione monoxime dictates transcriptional activation of ion channels and Ca(2+)-handling proteins. Mol. Pharmacol. 66, 708–17 (2004).

12. Ferreira, G., Artigas, P., Pizarro, G. & Brum, G. Butanedione Monoxime Promotes Voltage-dependent Inactivation of L-Type Calcium Channels in Heart. Effects on Gating Currents. J. Mol. Cell. Cardiol. 29, 777–787 (1997).

13. Tripathy, A., Xu, L., Pasek, D. A. & Meissner, G. Effects of 2,3-butanedione 2-monoxime on Ca2+ release channels (ryanodine receptors) of cardiac and skeletal muscle. J. Membr. Biol. 169, 189–98 (1999).

14. Hebisch, S., Bischoff, E. & Soboll, S. Influence of 2,3-butanedione monoxime on heart energy metabolism. Basic Res. Cardiol. 88, 566–575 (1993).

15. Cooling, M. T., Hunter, P. & Crampin, E. J. Sensitivity of NFAT Cycling to Cytosolic Calcium Concentration: Implications for Hypertrophic Signals in Cardiac Myocytes. Biophys. J. 96, 2095–2104 (2009).

16. Readnower, R. D., Brainard, R. E., Hill, B. G. & Jones, S. P. Standardized bioenergetic profiling of adult mouse cardiomyocytes. Physiol. Genomics 44, 1208–1213 (2012).

17. Ribeiro, A. J. S. et al. Multi-Imaging Method to Assay the Contractile Mechanical Output of Micropatterned Human iPSC-Derived Cardiac Myocytes. Circ. Res. 120, 1572–1583 (2017).

18. Iribe, G., Helmes, M. & Kohl, P. Force-length relations in isolated intact cardiomyocytes subjected to dynamic changes in mechanical load. Am. J. Physiol. Circ. Physiol. 292, H1487–H1497 (2007).

19. Janssen, P. M. L. & Periasamy, M. Determinants of frequency-dependent contraction and relaxation of mammalian myocardium. J. Mol. Cell. Cardiol. 43, 523–531 (2007).

20. Lee, S.-H. et al. REEP5 depletion causes sarco(endo)plasmic reticulum vacuolization and cardiac functional defects. Nat. Commun. in revisio, NCOMMS-18233B-Z

21. Sakurai, T. et al. Live Cell Imaging of Primary Rat Neonatal Cardiomyocytes Following Adenoviral and Lentiviral Transduction Using Confocal Spinning Disk Microscopy. J. Vis. Exp. e51666 (2014). doi:10.3791/51666

22. Blaser, M. C. et al. Deficiency of Natriuretic Peptide Receptor 2 Promotes Bicuspid Aortic Valves, Aortic Valve Disease, Left Ventricular Dysfunction, and Ascending Aortic Dilatations in MiceNovelty and Significance. Circ. Res. 122, 405–416 (2018).

23. Tseng, Q. et al. Spatial organization of the extracellular matrix regulates cell-cell junction positioning. Proc. Natl. Acad. Sci. 109, 1506–1511 (2012).

24. Hadipour-Lakmehsari, S. et al. Nanoscale reorganization of sarcoplasmic reticulum in pressure-overload cardiac hypertrophy visualized by dSTORM. Sci. Rep. 9, 7867 (2019).

